# Mechanisms of extracellular electron transfer in anaerobic methanotrophic archaea

**DOI:** 10.1101/2023.07.24.550278

**Authors:** Heleen T Ouboter, Rob Mesman, Tom Sleutels, Jelle Postma, Martijn Wissink, Mike S M Jetten, Annemiek ter Heijne, Tom Berben, Cornelia U Welte

## Abstract

Anaerobic methanotrophic (ANME) archaea are environmentally important uncultivated microorganisms mitigating the release of the potent greenhouse gas methane. During methane oxidation ANME archaea engage in extracellular electron transfer (EET) with other microorganisms, metal oxides, and electrodes, through a currently unknown mechanism. To shed light on this mechanism, we cultivated ANME-2d archaea (’*Ca*. Methanoperedens’) in bioelectrochemical systems and observed strong methane-dependent current (91-93% of total current) associated with high enrichment of ‘*Ca*. Methanoperedens’ on the anode (up to 82% of the community) determined by metagenomics and transmission electron microscopy. Electrochemistry and metatranscriptomics indicated that the EET mechanism was similar at various electrode potentials pointing to the involvement of an so far uncharacterized short-range electron transport protein complex and OmcZ nanowires, suggesting a unique EET pathway in all ANME-2 archaea. Our findings furthermore indicate that bioelectrochemical cells might be powerful tools for the cultivation, and possibly isolation, of uncultured electroactive microorganisms.

## Introduction

The anaerobic oxidation of methane is an important microbial process that regulates the release of methane, a potent greenhouse gas, into the atmosphere. This process is performed by anaerobic methanotrophic ANME-1, -2abc and -3 in marine sediments^1–5^, and by ANME-2d in freshwater sediments^6–10^. Our knowledge of the physiology of ANME is heavily reliant on -omics techniques as none of the ANME have been cultured independently of other microorganisms. Marine ANME are dependent on sulfate-reducing bacteria as an electron sink^11–13^. Freshwater ANME can couple methane oxidation to the reduction of nitrate to nitrite, a toxic byproduct. ANME-2d are therefore often found in consortia with nitrite-scavenging bacteria^6,7,14^. Alternatively, ANME can use insoluble metal oxides as electron acceptors, which likely makes these archaea an important methane sink in iron-rich sediments^15–19^

ANME-2d, which belong to the *Methanoperedenaceae* family, can use humic substances, manganese oxides, iron oxides, selenate, arsenate, and chromate as the electron acceptor for the conversion of methane^20–27^. For electroactive microorganisms the challenge is to adapt their protein machinery to tune into the different surface redox potentials of insoluble electron acceptors such as metal oxides, which are omnipresent in sediments^28^. Although the theoretical free energy gain increases the more positive the electron acceptor redox potential couple is, the enzymatic machinery of the microorganisms eventually determines whether this increased theoretical yield can be capitalized on^28^. Previous research has demonstrated that the extracellular electron transfer (EET) machineries of the model electroactive microorganisms *Geobacter* and *Shewanella* involve various extracellular electron transfer proteins, many of which contain multi-heme *c*-type cytochromes (MHCs) that form electron conduits. *Geobacter* encodes triheme periplasmic-type cytochromes (Ppc)^29,30^, pili^31^ and several outer-membrane cytochromes (OMCs) as part of its EET machinery^29,32^. Similar to these electroactive bacteria, ‘*Ca.* Methanoperedens’ species have a large MHC repertoire, some of which are expressed during growth with iron oxides, manganese oxides, or nitrate as the electron acceptor^20,24,33^. However, the mechanism of extracellular electron transfer in ANME archaea remains largely unresolved. While some MHCs have been implicated in EET, comparative physiology studies of ANME archaea are scarce due to the limitations imposed by their slow growth rates in complex communities, and cryptic cultivation requirements. These problems are compounded by the presence of bacteria known to perform extracellular electron transfer, such as *Geobacter,* in *Methanoperedenaceae*-dominated enrichment cultures. It has been suggested that soluble intermediates produced by ANME archaea (i.e., acetate) are utilized by other electroactive bacteria, which then perform the ultimate reduction of the extracellular electron acceptor^34–36^. In this study, we aimed to investigate extracellular electron transfer by ‘*Ca*. Methanoperedens’ ANMEs using bioelectrochemical systems at various anode potentials to perform comparative physiology experiments. Here we present the resulting bioelectrochemical data, visualization of the anode biofilm using fluorescence and electron microscopy, and metagenomics and metatranscriptomics analyses of the composition and activity of said biofilm. We reached a strong methane-dependent current that was not dependent on the anode potential. Analysis of gene expression showed upregulation of two gene clusters possibly involved in EET. At the same time, we obtained highly enriched microbial communities dominated by ‘*Ca*. Methanoperedens’ that might open doors to future axenic ANME cultures.

## Results

In total, three experiments were performed to investigate the mechanism of extracellular electron transfer (EET) by *’Ca*. Methanoperedens to a gold anodic electrode mesh by several tests and sampling for metagenomic and metatranscriptomic analysis and microscopy imaging (Figure 1). In **experiment 1** (Figure S1), there was a start-up phase of approximately one week where four parallel systems were run at 0 mV vs SHE to develop a comparable biofilm and a subsequent phase of two more weeks where the potential was kept at 0 mV in BES1 or switched to 200 mV (BES2), 400 mV (BES3), or 600 mV (BES4). In **experiment 2** (Figure **S*2***), the microbial community was incubated for 6.5 weeks at 0 mV vs SHE in a BES. In **experiment 3** (Figure S3), the microbial community was incubated for 9 weeks at 0 mV vs SHE in a BES.

**Figure 1:**
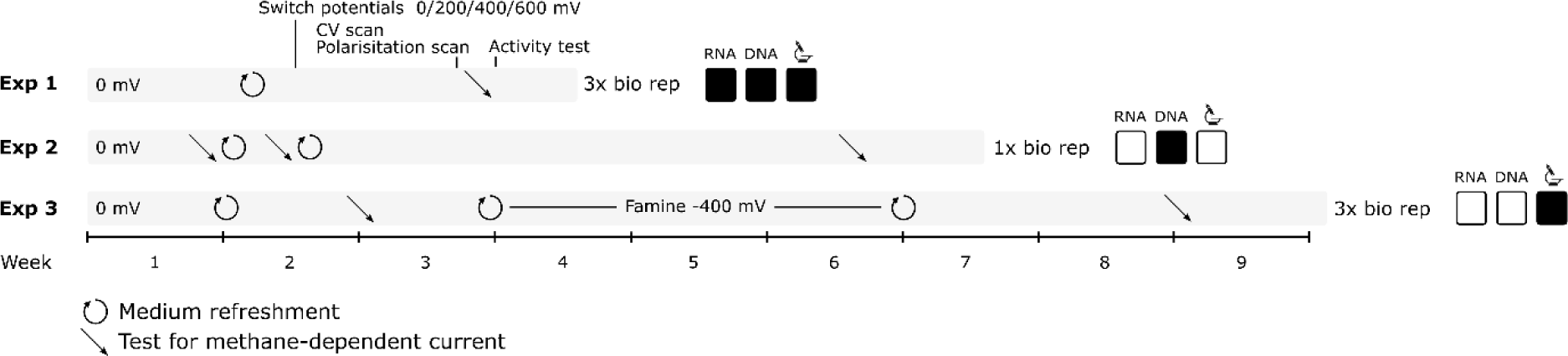
A schematic overview of the three performed experiments. Events during the experiments such as medium refreshments, test for methane-dependent current, cyclic voltammetry (CV) and polarisation scans, activity test, and sampling for metagenomics and metatranscriptomics analysis as well as microscopy visualization are indicated.

### Methane-dependent current production remains unchanged at varying anode potentials

To investigate the mechanism of EET by *’Ca*. Methanoperedens’ and the influence of the electric potential on the process, we conducted bioelectrochemical experiments at four different anode potentials: 0 mV, 200 mV, 400 mV, and 600 mV vs SHE (Figure S1). In each BES, the initial biofilm was established under near-identical conditions at a potential of 0 mV vs SHE, after which the potential was changed, and current production and the microbial community were monitored (**experiment 1**). No clear effect of the poised potential on the amount of produced current was observed (Figure 2A), while a strong methane-dependent current (59% ± 11% out of 39 ± 8.8 mA m^-2^) was produced at all potentials (Figure 2B, Figure S4). Anaerobic methanotrophic activity was confirmed through the increase of ^13^CO_2_ over total CO_2_ after the culture was supplied with ^13^CH_4_ (Figure S4). The microbial community consumed on average 71 ± 0.017 μmol methane day^-1^ which was accompanied by the production of 3.0 ± 1.5 C (Figure S4). Cyclic voltammetry scans at different anode potentials were nearly identical, suggesting that the same redox centres were operational under the four different poised anode conditions (Figure 2C). Polarization curves indicated that the potential at which the microbial community began to produce current was close to the thermodynamic redox potential of the CH_4_/CO_2_ redox couple at -0.249 V (with conditions: pH 7.25 and a gas phase of 86.4% CH_4_ and 4.55% CO_2_) (Figure 2D). The shape of the polarization curves was similar, suggesting that the EET mechanism was not influenced by the applied potential (Figure 2D).

**Figure 2:**
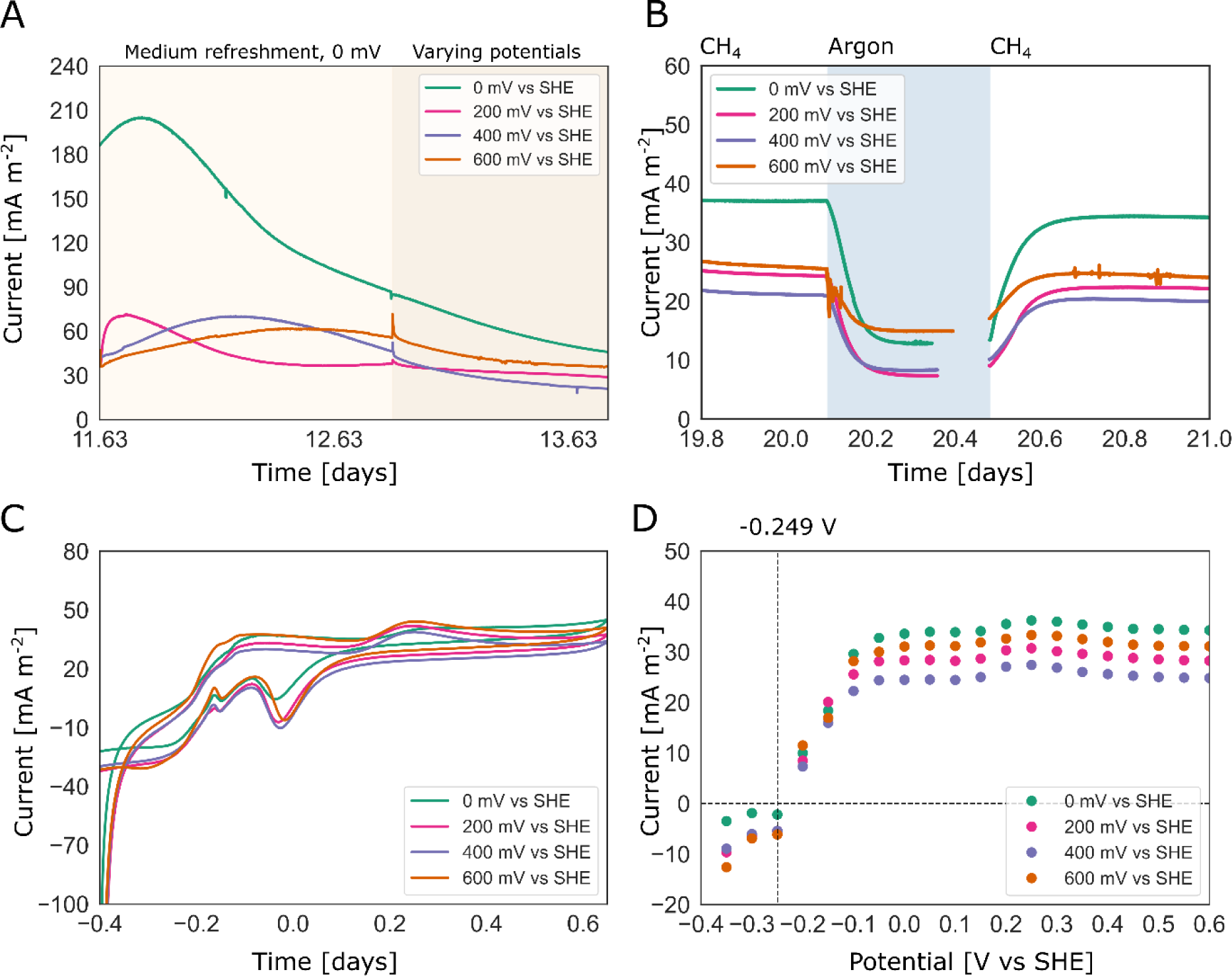
Bioelectrochemical data obtained during experiment 1, including measurements taken under various potentials. The figures show the average of three biological replicates. A) The current production after the medium was replaced and after all systems were initially operated at 0 mV vs SHE, followed by a switch to different potentials (0 mV, 200 mV, 400 mV, or 600 mV vs SHE) B) The replacement of methane by argon to test for methane-dependent current, C) Cyclic voltammetry scans, D) Polarisation scans with the redox potential of the CH_4_/CO_2_ redox couple indicated by the dashed line at -0.249 V

### Long-term incubations in bioelectrochemical systems led to anode biofilms highly enriched in ‘*Ca*. Methanoperedens’

After having established stable methane-dependent current production at 0 V in **experiment 1** (Figure S1), we prolonged the incubation time, performed additional medium refreshments, and monitored the percentage of methane-dependent current in **experiment 2**. The relative abundance of ‘*Ca.* Methanoperedens’ was determined at the end of each incubation (**experiments 1, 2**; Figure 3 and Figure S*5*), as sampling of the electrode biofilm requires sacrificing the experiment. We conducted separate experiments with different incubation times and medium refreshments (**experiment 1** and Figure S1; **experiment 2** and Figure **S*2***). Our results showed that a higher number of medium refreshments and a longer incubation time were associated with higher percentages (up to 91%) of methane-dependent current and higher relative abundance (up to 82%) of *’Ca*. Methanoperedens’ at the anode (Figure 3 & Figure S*5*) determined by metagenomics.

**Figure 3:**
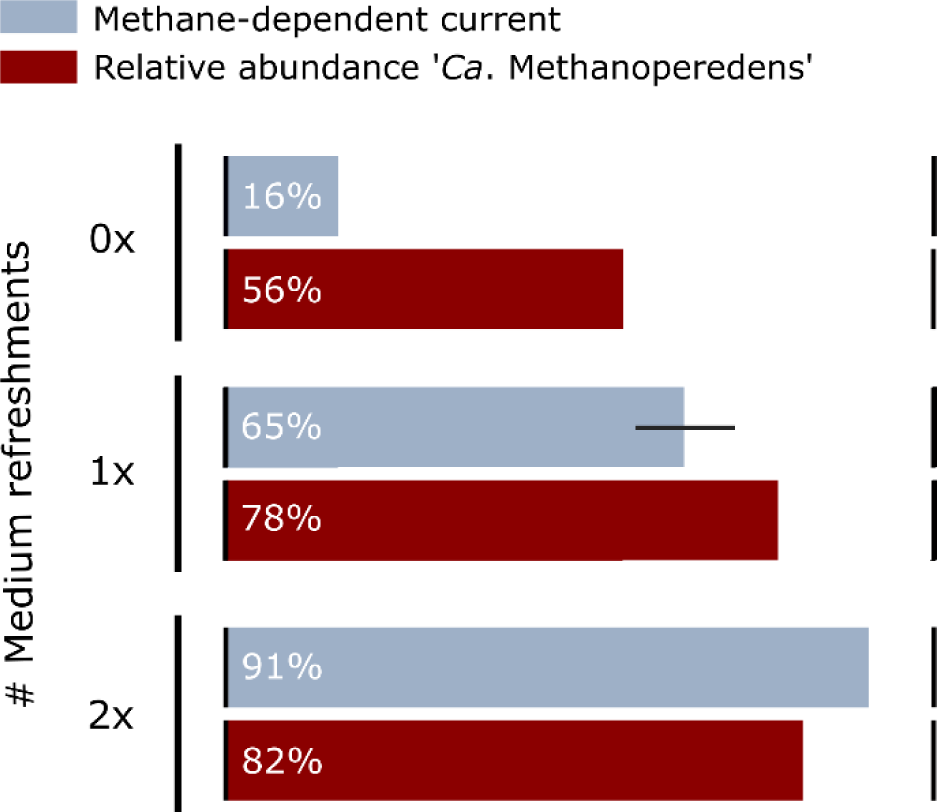
**Influence of the number of medium refreshments and incubation time on the percentage of methane-dependent current and relative abundance of ‘*Ca*. Methanoperedens’.**

In **experiment 3**, during three out of nine weeks we applied a potential of -400 mV vs SHE (Figure S3). By applying a potential that is lower than -0.24 mV, which is the potential of redox couple CH_4_/CO_2_, we tested whether anaerobic methane oxidation can be reversed by ‘*Ca.* Methanoperedens’. This period turned out to be a starvation period as we observed that the culture could only produce little methane (0.22 ± 0.029 μmol methane day^-1^) and a stable current of -0.08 mA m^-2^ which shows that they can only reach at best a methane production rate that is 0.31% of the methane oxidation rate. Nonetheless, the microbes remained active even after the starvation period with 41 mA m^-2^ of stable current before starvation vs. 61 mA m^-2^ after starvation (Figure S3). After starvation, when the potential was returned to 0 mV vs SHE, 93% methane-dependent current was produced (Figure S*5*E, Figure S3).

### Imaging revealed spatial organization of the biofilm microbial community

Confocal laser scanning microscopy combined with fluorescence in-situ hybridization (FISH), scanning electron microscopy (SEM), and transmission electron microscopy (TEM) were employed for samples from **experiment 3** to visualize the biofilm on the anode (Figure 4). The signals for ‘*Ca.* Methanoperedens’ and archaea fully overlapped indicating that no other archaea than ‘*Ca.* Methanoperedens’ were present (data not shown), congruent with the meta-omics data (Figure 5). The confocal laser scanning micrographs revealed a large number of *’Ca*. Methanoperedens’ cell aggregates (Figure 4B, F); the fluorescent signals of ‘*Ca*. Methanoperedens’ aggregates, together with the laser reflection used to visualize gold, were both observable in a confocal volume of 500 nm, governed by the system’s optical resolution determined by the used 63x/1.40 lens. This is evidence that ‘*Ca*. Methanoperedens’ are present further than 500 nm away from the anode gold mesh (Figure 4B) and that they were also present within 500 nm of the gold mesh (Figure 4F). Since *Geobacter* sp. is a well-known electro-active bacterium hypothesized to be involved in EET by *’Ca*.

**Figure 4:**
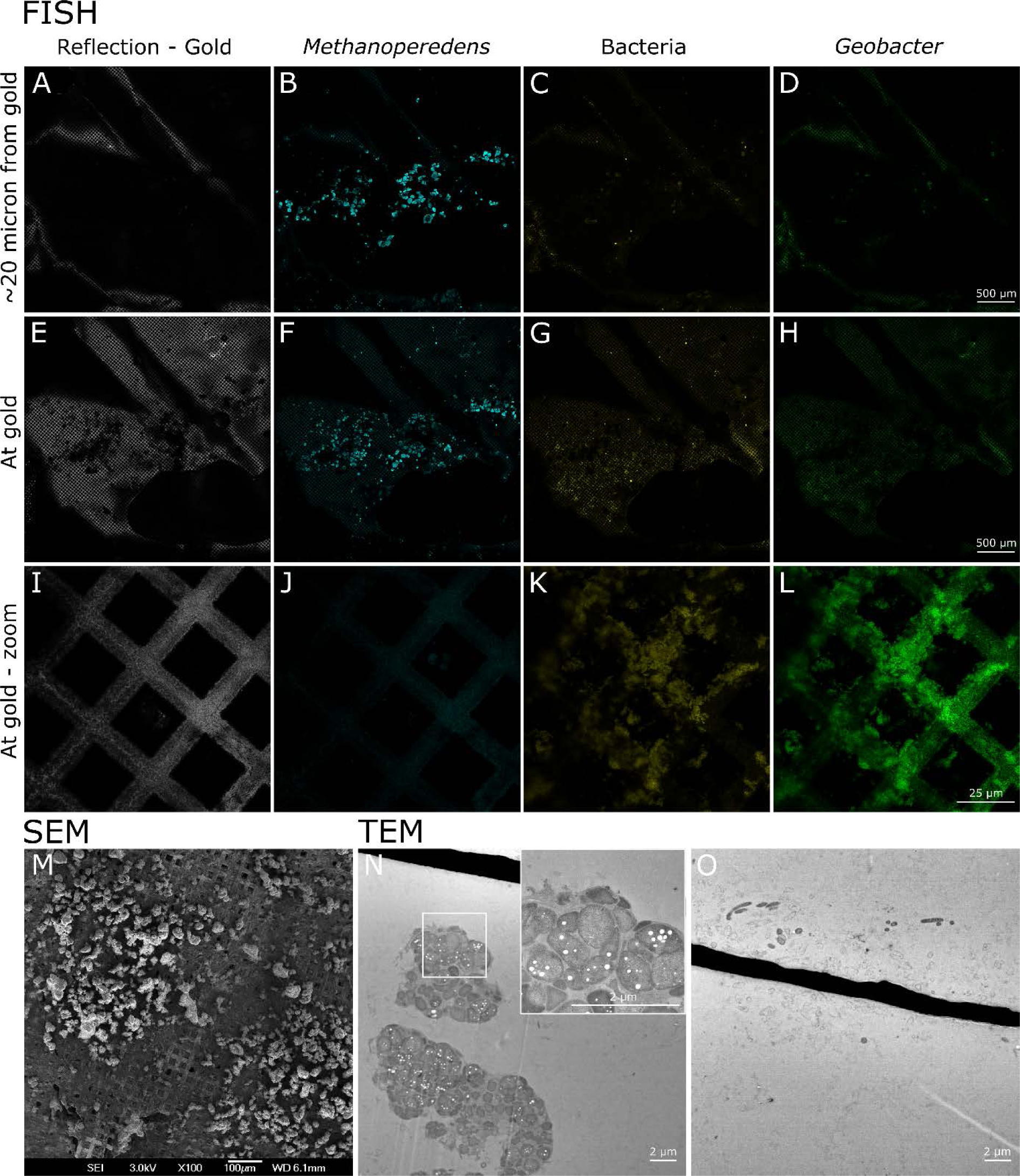
Visualization of the biofilm using different microscopy techniques. A-L: Confocal laser scanning microscopy combined with fluorescence *in situ* hybridization (FISH) to target *’Ca*. Methanoperedens’ using a FLUOS probe (cyan), bacteria using a Cy5 probe (yellow), and *Geobacter* sp. using a Cy3 probe (green), M, N: transmission electron microscopy (TEM) indicating live *Ca*. Methanoperedens cells (M) further away from the gold mesh and rod-shaped bacterial cells (N) closer to the gold mesh, many of which appeared to be empty and therefore dead, O: scanning electron microscopy (SEM) showing large clusters resembling ‘*Ca*. Methanoperedens’ cells further away from the electrode mesh and a dense biofilm of rod-shaped bacteria closer to the gold mesh. All the samples shown in this figure were obtained from experiment 3.

**Figure 5:**
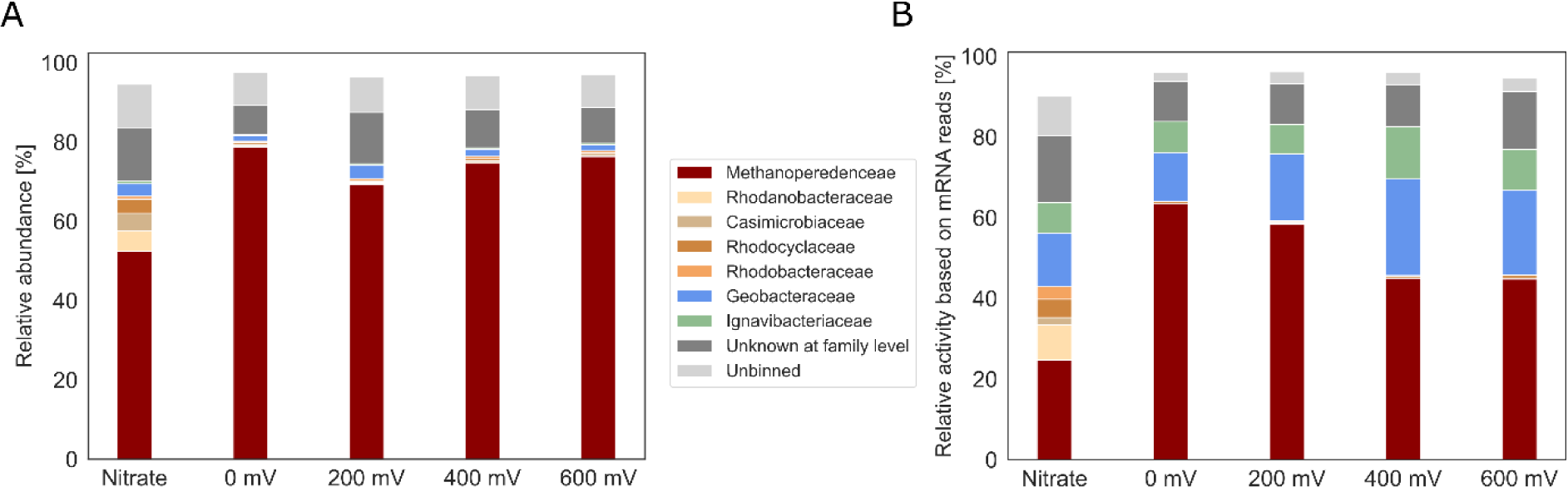
Relative abundance and relative activity at the family level determined by mapping (A) metagenome reads and (B) RNA-seq reads to the metagenome-assembled genomes (MAGs). The families *Methanoperedenaceae* and *Geobacteraceae* both comprise a single MAG that was identified as ‘*Ca*. Methanoperedens’ and *Geobacter* sp., respectively, based on the classification by the Genome Taxonomy Database (GTDB). Only taxa with a relative abundance or activity of greater than 2.5% are displayed.

Methanoperedens’ and part of the microbial community in the inoculum^37^, we targeted this bacterium specifically. Bacteria were observed near the mesh, and a large proportion of these bacteria hybridized with the *Geobacter* probe (Figure 4K, L). Scanning electron microscopy revealed granules with the typical cell morphology of ‘*Ca*. Methanoperedens’, which covered a large portion of the electrode (Figure 4M). Correspondingly, transmission electron microscopy showed aggregated sarcina-shaped clusters, with intact cells containing storage compounds that were represented as white dots within the granule (Figure 4N). The clusters were located at varying distances from the gold electrode mesh with a minimum of 4 μm (Figure 4N), corresponding with the FISH observations, and a maximum of 45 µm (data not shown). Furthermore, the transmission electron micrographs showed bacterial cells in the proximity of the gold mesh, many of which appeared to be dead cells, based on the loss of structural integrity and the absence of cell content (Figure 4O).

### Antibiotics targeting bacteria do not affect current production

To unravel the relative contribution of bacteria and archaea to the current production, we incubated the anode biofilms with antibiotics, consisting of vancomycin, streptomycin, ampicillin and kanamycin, targeting only bacteria and not archaea. The current did not change after the addition of antibiotics, with a small increase of current observed right after the antibiotic addition (Figure S7, Figure S8, Figure S9). After several days of incubation, the current did not drop below the current density measured before the addition of antibiotics and the methane-dependency of the current increased by 2%, corroborating the negligible role of bacteria in current generation. To further test the hypothesis that ‘*Ca*. Methanoperedens’ is the principal driver of current generation we added 2- bromoethanosulfonate, which inhibits the key methanotrophy enzyme methyl-coenzyme M reductase (MCR), and puromycin, an antibiotic that affects the RNA translation with a more pronounced effect on archaea compared to bacteria. The addition of 20 mM 2- bromoethanosulfonate resulted in an immediate and strong reduction in current by 89% (Figure S9), suggesting that a small proportion of the current may be generated by the conversion of storage polymers, such as polyhydroxyalkanoate (PHA), which were visible in the TEM micrograph (Figure 4N), a process that is independent of MCR. Upon the addition of puromycin, the current showed a gradual decrease, indicating that the antibiotics successfully penetrated the biofilm and affected the archaea (Figure S8); at the same time, it establishes that the remaining current was due to residual activity of enzymes in puromycin-affected archaeal cells, unlike what was seen in the treatment with bacterial-targeting antibiotics, where no gradual decrease of current density was observed (Figure S8 and Figure Figure S*9*).

### Global metagenome & metatranscriptome analysis

Metagenomics and metatranscriptomics analyses were conducted to further determine the composition of the microbial community and the activity of its members (Figure 5,Figure S8). The relative abundance of ‘*Ca*. Methanoperedens’ was 54% under nitrate reducing conditions, and reached a maximum of 78% in the anode biofilm after three weeks of incubation (Figure *5*) and 82% after 9 weeks of incubation (Figure 3 and FigureFigure S3). The transcriptional activity was measured as the amount of RNA-seq reads mapping to a specific genome relative to the number of reads mapping to the total metagenome. ‘*Ca*. Methanoperedens’ represented approximately 20% of the total activity under nitrate reducing conditions, whereas it represented 45-64% of the total activity in the BES when the electrode was used as an electron acceptor (Figure 5B, Figure S10). Several proteobacteria were found to be active in the nitrate condition which were not active in the BES. *Geobacteraceae* and *Ignavibacteriaceae*, despite their low relative abundance, were found to be active under all conditions. These bacteria are known for their electroactive properties and were probably using organic matter derived from decaying biomass or excreted products as their electron donor. The completeness of the ‘*Ca*. Methanoperedens’, *Geobacter* sp. and *Ignavibacteriaceae* MAGs were high with 99.3%, 99.4% and 97.2% respectively.

### Transcriptomics analyses reveal two differentially expressed gene clusters most likely involved in extracellular electron transfer

The ‘*Ca*. Methanoperedens’ MAG contains genes for the complete reverse methanogenesis pathway In addition to several more copies detected in the unbinned contigs of the metagenome. Although several strains of ‘*Ca*. Methanoperedens’ were present in the BES, only one was highly abundant and transcriptionally active (Table S2A). The genes within the MAG encoding proteins of the reverse methanogenesis were upregulated in the electrode condition compared to the nitrate-reducing condition indicating the high activity of ‘*Ca.* Methanoperedens’. Using metatranscriptomics, we identified two strongly upregulated gene clusters in this MAG that encoded multiple multi-heme *c*- type cytochromes (MHCs). The first gene cluster contained two 7-heme binding MHCs (Metp_01715, Metp_01720) and was co-located with genes encoding a NrfD-like *b*-type cytochrome-containing transmembrane protein (Metp_01713), a protein predicted to harbor four [4Fe-4S] clusters (Metp_01714), an Fdh-like membrane-integral *b*-type cytochrome (Metp_01719), and unknown proteins (Metp_01716-18, Metp_01721-22) (Figure *6*). While some genes in this cluster were expressed under both electrode and nitrate conditions, others were upregulated in the electrode condition (Table S2A, Figure *6*). The second gene cluster contained genes that encode proteins homologous to those found in *Geobacter sulfurreducens* related to the formation of OmcZ nanowires^38^: Metp_02468 encodes a protein that is homologous to OmcZ from *Geobacter sulfurreducens* and Metp_02469 encodes a protein that is homologous to a serine protease OzpA from *G. sulfurreducens* that is required for the assembly and folding of the OmcZ extracellular nanowire.

**Figure 6:**
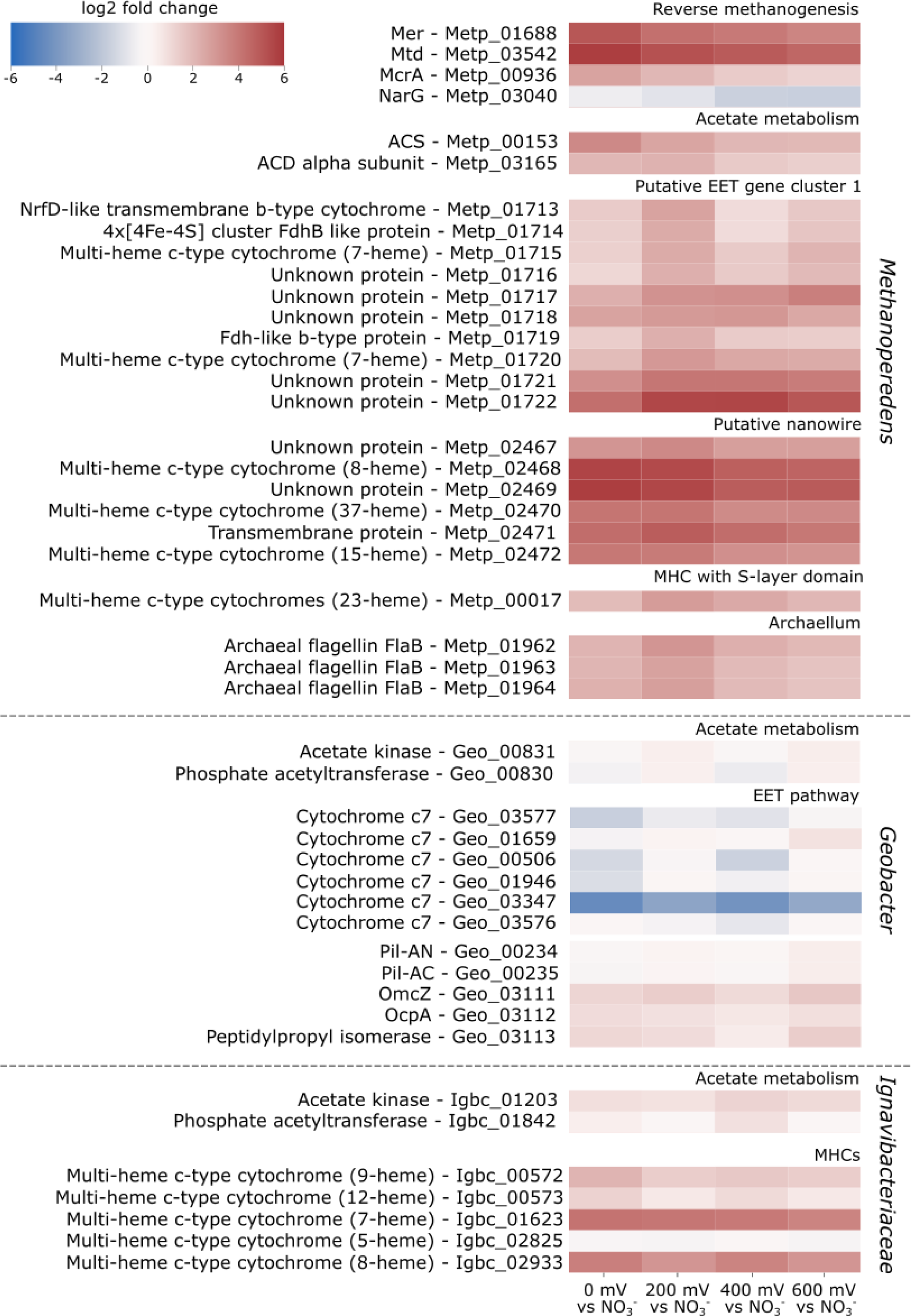
Differential gene expression analysis comparing the electrode condition with potentials 0 mV, 200 mV, 400 mV or 600 mV to the nitrate condition. The Figure displays two gene clusters from ‘*Ca*. Methanoperedens’ that are potentially involved in EET, along with genes involved in the EET mechanism of *Geobacter* sp. Additionally, it shows MHCs from *Ignavibacteriaceae* with at least thirty normalized counts in the electrode condition, as the EET mechanism of this microorganism is currently unknown. *Geobacter* and *Ignavibacteriaceae* were specifically mentioned as they were found to be active in the study. Furthermore, genes encoding enzymes involved in acetate metabolism from these three microorganisms were added, as they may play a role in the interaction between ‘*Ca*. Methanoperedens’ and the two bacteria.

All genes in this cluster were strongly expressed (Table S2A) and upregulated in the electrode condition, with log2-fold changes of at least 3.6 (Figure *6*). A schematic overview of the metabolism of ‘*Ca.* Methanoperedens’ and its proposed EET mechanism is depicted in Figure *7*.

**Figure 7:**
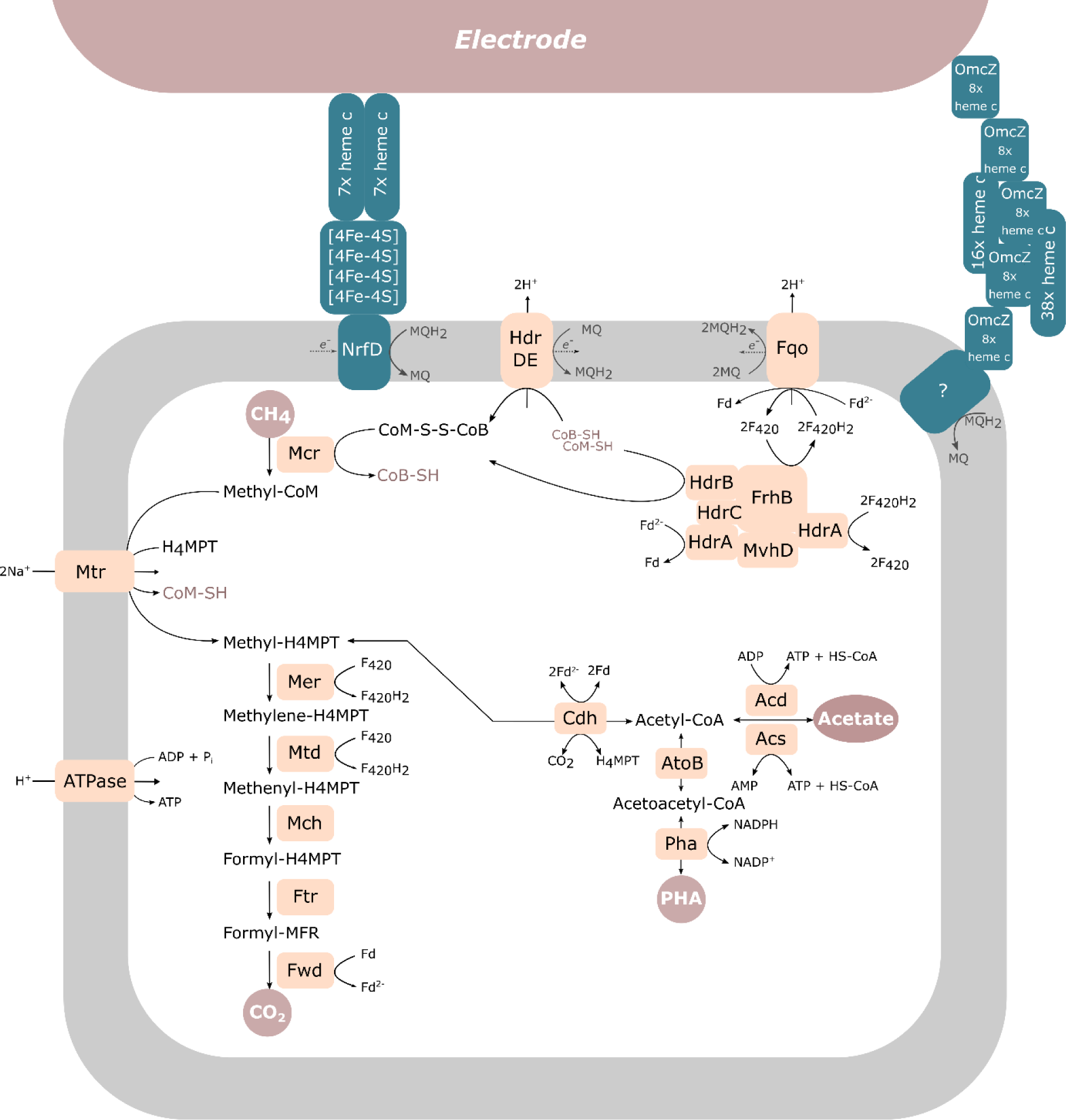
**Overview of metabolism from ‘*Ca.* Methanoperedens’ and its putative EET mechanism towards the electrode.**

We also mined the *‘Ca.* Methanoperedens’ MAG for multi-heme *c*-type cytochromes and found fifteen out of a total of thirty-three MHCs upregulated (log2-fold change > 2) at the electrode compared to the nitrate condition (Table S2B). One of these MHCs had an S-layer domain suggesting its involvement in the electron transfer through this outer layer. Additionally, all three genes encoding major subunit flagellin (*flaB*) that were found in the *Methanoperedens* MAG were upregulated in the electrode condition. *Geobacter* and *Ignavibacteriaceae* were both active at the electrode (Figure *6*): *Geobacter* expressed genes related to their EET mechanism together with genes related to acetate metabolism (Table S2A), but none of these genes was significantly upregulated (log2-fold change > 2) at the electrode compared to the nitrate condition. For *Ignavibacteriaceae* the EET mechanism is not known but we found upregulated genes encoding MHCs (Figure *6*, showing *Ignavibacteriaceae* MHCs > 30 normalized counts). Genes related to their acetate metabolism were expressed but not upregulated (Table S2C).

The metatranscriptome analysis indicated that the expression patterns observed for the four different potentials were similar, as previously observed in the bioelectrochemical data (Figure 2,Figure *6*).

## Discussion

In this work, we demonstrated the ability of ‘*Ca.* Methanoperedens’ to perform extracellular electron transfer (EET) to an electrode at potentials ranging from 0 mV to 600 mV vs standard hydrogen electrode (SHE). We discovered that during this process, ‘*Ca*. Methanoperedens’ expresses a gene cluster encoding for proteins with structural similarities to the OmcZ nanowire-related proteins found in *Geobacter sulfurreducens* used for long-range (100 µm) electron transfer with the highest expression at 0 mV^38^. Unlike the OmcZ nanowire found in *Geobacter*, *’Ca*. Methanoperedens’ encodes two additional multiheme *c*-type cytochromes (MHCs) with 37 and 15 heme-binding sites, respectively, in the same gene cluster. We identified that this gene cluster was highly upregulated (log2-fold change > 3.5) under electrogenic growth conditions compared to when nitrate was used as an electron acceptor (Table S2A). Surprisingly, varying the potential from 0 to 600 mV vs. SHE did not influence the EET mechanism of ‘*Ca.* Methanoperedens’. This was evidenced by the presence of similar redox centers across the different tested conditions, the comparable shape of the polarization scans and the absence of potential-induced changes in current production and expression patterns (Figure 2, Figure *6*, Table S2). This suggests that ‘*Ca.* Methanoperedens’ may be using diverse extracellular electron acceptors in the environment without the need to change its expression pattern or EET mechanism, unlike what was reported for *Geobacter* which can fine tune its EET mechanism in response to a presented redox potential^39–41^. *Geobacter’*s requirement for such adaptation may be attributed to the significant number of heterotrophic competitors it encounters, whereas for *’Ca*. Methanoperedens’, this number is probably considerably smaller, given its status as a chemolithotrophic autotroph.

Although EET by ‘*Ca.* Methanoperedens’ has been studied both in bioelectrochemical systems and in relation to metal oxides, in most studies the EET mechanism has not further been unraveled^42–47^. Two exceptions are studies focusing on the use of ferrihydrite and birnessite by ‘*Ca.* Methanoperedens’, which identified several MHCs expressed during metal-oxide reduction and resulted in the classification of three new *Methanoperedenaceae* members ‘*Ca.* Methanoperedens ferrireducens’, ‘*Ca.* Methanoperedens manganicus’ and ‘*Ca.* Methanoperedens manganireducens’^20,24^. Similar to ‘*Ca.* Methanoperedens manganicus’ the *Methanoperedenaceae* member in this study expressed genes encoding a large subunit flagellin (*flaB*) that is part of the archaellum which has been found to be electrically conductive in *Methanospirillum hungatei*^48^. Studies investigating the role of MHCs containing an S-layer domain in extracellular electron transfer have reported varying results depending on the electron acceptor used. Leu *et al.* have shown the expression of such MHCs during manganese-dependent methanotrophy^24^, while no such MHCs were expressed during EET to iron minerals or syntrophic growth with sulfate-reducing bacteria^20,49^. We found one MHC with 23 heme-binding motifs in our ‘*Ca.* Methanoperedens’ MAG fused with a putative S-layer domain as described by McGlynn *et al.,* 2015 ^13^; the gene was differentially expressed during electrogenic growth compared to nitrate-dependent growth. The presence of this MHC with fused S-layer domain suggests its involvement in electron transfer across the outer layer of the archaeon while using the electrode as electron acceptor.

The expression of a second gene cluster containing genes encoding for MHCs, an [4Fe-4S] cluster protein and *b*-type cytochromes was observed during both electrode utilization and the use of nitrate as an electron acceptor. While it was strongly upregulated under growth on electrodes with the highest expression at 200 mV, it had a high baseline expression under nitrate-dependent growth conditions (with an average of 202 normalized counts for the genes in this cluster) compared to genes involved in reverse methanogenesis (*mer,* 39; *mch,* 6; *mtd,* 15). Given these findings, this gene cluster may be involved in a process unrelated to the reduction of solid electron acceptors, such as direct interspecies electron transfer (DIET), like the process observed between *Geobacter* and methanogens^50–53^, or short-range electron transfer to extracellular electron acceptors. Interestingly, Leu *et al*. identified a similar gene cluster in ‘*Ca*. Methanoperedens manganicus’ that was found to be highly expressed during growth with birnessite, making its involvement in extracellular electron transfer likely.

The inaccessibility of a pure culture of any of the sub-clades of anaerobic methanotrophic (ANME) archaea has created a significant knowledge gap regarding their physiology. ANME archaea are reliant on syntrophic partners to act as electron sinks^11–13^ or to convert nitrite^6,14,54^, which is a toxic byproduct of the independent reaction ANME-2d can perform converting methane at the expense of nitrate. In this study, we successfully overcame this obstacle by cultivating ANME-2d on an electrode. We obtained a culture where *’Ca*. Methanoperedens’ comprised 82% of the total community, which was achieved in a timeframe of only a couple of weeks. This was accompanied by a produced current that was 93% directly dependent on methane, suggesting that bioelectrochemical systems might be key in further enriching and possibly eventually obtaining axenic ANME cultures. Although *Geobacter* and *Ignavibacteriaceae* were active at the electrode, the lack of effect of bacterial antibiotics streptomycin, vancomycin, ampicillin and kanamycin on the methane-dependent current and the significant impact of 2-bromoethanosulfonate, an MCR inhibitor, on the current indicates the low contribution of these bacteria to the total current produced. As *Geobacter* sp. is known to be inhibited by streptomycin and kanamycin^55^, this excludes the possibility that the methane-dependent current is produced via *Geobacter* through direct interspecies electron transfer^34–36^. Additionally, the intact ‘*Ca*. Methanoperedens’ clusters contrasting the dead bacterial cells visualized using various microscopic techniques provide further evidence for ‘*Ca.* Methanoperedens’ to be the driving microorganism for the produced current and confirms that ‘*Ca.* Methanoperedens’ can perform electrogenic anaerobic oxidation of methane independently of syntrophic partners.

In conclusion, we present evidence for direct current production and extracellular electron transfer by ‘*Ca*. Methanoperedens’ during methane oxidation without the need of syntrophic partner microorganisms, possibly via cytochrome-containing nanowires. These findings have implications for anaerobic methanotrophs in the environment as most forms of methanotrophy rely on extracellular electron transfer; these methanotrophs together represent the major anoxic methane sink on our planet with strong implications for climate change. At the same time, direct electricity production from methane offers exciting opportunities for future sustainable biotechnological applications.

## Methods

### Cultivation of ‘*Ca.* Methanoperedens’ in bioelectrochemical systems

The inoculum was obtained from an enrichment culture dominated by ‘*Candidatus* Methanoperedens’ seeded from Vercelli rice fields that was exposed to nitrate limiting conditions and continuously operated since March 2014 ^37,54^. This culture was grown on electrodes in a bioelectrochemical system (BES) as previously described^33^ with a few adaptations. The BES medium consisted of (per L) 0.1 g CaCl_2_·2H_2_O, 0.1 g MgCl_2_·4H_2_O, 0.05 g KH_2_PO_4_, 0.5 g NH_4_Cl, 2.38 g HEPES and was autoclaved prior to the addition of 2 mL L^-1^ trace elements and 0.1 mL vitamin solution with final pH of 7.25 and room temperature. After inoculation, 0.3 mL FeSO_4_ (10 mM) was added to the anaerobic anolyte and the anode chamber was continuously sparged with CH_4_/CO_2_ 95%/5% at 10 mL min^-1^ and N_2_ at 2.2 mL min^-1^ while the cathode chamber was continuously sparged with N_2_ at 10 mL min^-1^. The two chambers from the BES were separated via a a RALEX^®^ heterogeneous cation-exchange membrane (CMHPES) (Mega, Prague, Czech Republic). The anode electrode (7.7×1.9 cm) consisted of gold mesh foil (Precision Eforming, New York, US). Three experiments were performed (Figure *1*): in **experiment 1**, the BES was operated in two stages for a total duration of 3 weeks. In stage 1 (7-8 days) the anode was poised at a potential of 0 V vs SHE using a MultiEmStat3 potentiostat (PalmSense, Houten, Netherlands) to develop a biofilm under comparable conditions. At the end of stage 1, the medium containing planktonic cells was replaced by fresh medium and the electrode potential was set at a variety of potentials: 0.0 V, 0.2 V, 0.4 V and 0.6 V vs SHE. Stage II of the experiment was run for 14 d. At the end of the experiment RNA, DNA and microscopy samples were collected anaerobically. To compare RNA expression between the microbial community grown at the electrode and with nitrate as electron acceptor, a batch experiment was performed with the medium as mentioned above but with the addition of 3 mM nitrate, without the addition of a poised electrode. This experiment was run for five days after which samples were collected anaerobically for DNA and RNA sequencing. In **experiment 2**, the BES was operated at 0 mV including two medium refreshments with a total duration of 6.5 weeks to investigate the effect of a longer incubation on the community and the obtained current. At the end of the experiment DNA samples were collected anaerobically for metagenomic sequencing. In **experiment 3,** the BES was operated for 9 weeks at 0 mV including three medium refreshments and a 3-week famine phase at a potential of -400 mV. At the end of the experiment, DNA and microscopy samples were collected anaerobically.

In all three experiments we tested for methane-dependent current in different stages (Figure *1*): the gas inflow of CH_4_/CO_2_ 95%/5% was stopped and replaced by Argon/CO_2_ 95%/5%. For **experiment 1**, polarisation curves and cyclic voltammetry scans were recorded (Figure *1*). In addition, the gas flow into and out of the BESs was interrupted and 20 mL ^13^CH_4_ was added to the anolyte while ^13^CO_2_ and total CH_4_ were measured as described in Ouboter *et al*., 2022^37^ to verify methanotrophic activity.

The three experiments were performed over a period of several months to evaluate the reproducibility of the outcome**Error! Reference source not found.**. Negative controls were performed in previous experiments with dead and without biomass ^37^.

In experiment 4, a mixture of bacterial antibiotics was added to examine the involvement of bacteria in the current production. This mixture consisted of vancomycin HCl (Merck Life Science, Amsterdam, NL), streptomycin sulfate, ampicillin sodium salt, and kanamycin sulfate (VWR, Amsterdam, NL) added at a concentration of 50 μg mL^-1^ per antibiotic. We tested for methane-dependent current before and after the addition of the antibiotics mixture. Additionally, we added 20mM 2- bromoethanosulfonate (Merck Life Science, Amsterdam, NL) to two of the biological replicates and 50 μg mL^-1^ puromycin dihydrochloride (Merck Life Science, Amsterdam, NL) to two other biological replicates.

### Nucleic acid extraction from ‘*Ca.* Methanoperedens’ enrichment culture

RNA samples were taken from experiment 1 and DNA samples were taken from experiment 1 and 2. DNA was isolated following the Powersoil DNeasy kit protocol with the addition of a 10 min bead beating step at 50 s^-1^ (Qiagen, Hilden, Germany). RNA was isolated following the Ribopure Bacteria kit protocol (Thermo Fisher Scientific, Waltham, US), with the addition of a 15 min bead beating step at 50 s^-1^. rRNA depletion was tested according to Phelps *et al.* 2020^56^ but this method was unsuccessful. Instead, the NEB microbe rRNA depletion protocol was used (Macrogen, Seoul, South Korea). The metatranscriptomic datasets were constructed from biological and technical replicates, investigating five different conditions: 0 mV (2 biological, 1 technical replicate), 200 mV (3 biological replicates), 400 mV (3 biological replicates, 1 technical replicate), 600 mV (3 biological replicates, 1 technical replicate), nitrate condition (3 biological replicates).

### Metagenomic and metatranscriptomic dataset generation

For metagenome sequencing, the library was prepared using the TruSeq DNA PCR-Free Kit (Illumina, San Diego, CA, United States) and sequenced using Illumina NovaSeq 6000, obtaining 100 M 300-bp paired-end reads (Macrogen, Seoul, South Korea). The obtained reads were processed according to Ouboter *et al.,* 2022. For the metatranscriptome sequencing, the library was prepared using the TruSeq Stranded Total RNA Kit (Illumina, San Diego, CA, United States) and sequenced using the Illumina NovaSeq 6000 obtaining 100 M 150-bp paired-end reads. Sequence trimming was performed using Sickle and the obtained reads were pseudo-aligned against the predicted genes of the metagenome using Kallisto v0.46.1, the outcome was represented as transcript per million (tpm) value and counts. The data was further processed by removing genes with less than one count per million and by normalizing the counts using a weighted trimmed mean of the log expression ratios (trimmed mean of M values (TMM)) removing the effect of biological differences between samples making it possible to look at differential expression of genes^57,58^. This method assumes that the majority of the genes is not differentially expressed^58^. All sequencing data are available in the European Nucleotide Archive under BioProject PRJNA995526.

### Visualisation of the biofilm

Gold electrodes were removed from the BES systems under anoxic conditions. Electrode biofilm was fixed for 1h anoxically using 4% paraformaldehyde, 100 mM MOPS pH 7.3 after which samples were stored in the fridge in 0.1% paraformaldehyde and 100 mM MOPS pH 7.3 until further use.

### FISH imaging with confocal laser scanning microscopy

Biomass was dehydrated incubating the samples for 3 min stepwise increasing the ethanol concentration from 25% ethanol until 100%. Pieces of gold electrode were positioned on microscope slides and samples were air-dried for 30 min. A barrier was created around the samples using Silicone grease. Hybridisation buffer containing 180 μL 5M NaCl, 20 μL 1M Tris/HCl pH 8.0, 1 μL 10% w/v SDS, 350 μL formamide, 450 μL miliQ water, was carefully added to the samples covering the gold foil in solution while keeping the gold foil attached to the slide. Fluorescence *in situ* hybridization (FISH) was performed using probes labelled with Cy3, Cy5 and FLUOS that were added to the hybridization buffer at 10% of the total volume: 5’-GGTCCCAAGCCTACCAGT-3’-FLUOS (targeting ‘*Ca.* Methanoperedens’) ^59^, 5’-GTGCTCCCCCGCCAATTCCT-3’-Cy3 (targeting archaea) ^60^, a mixture of three probes 5’-GCTGCCTCCCGTAGGAGT-3’-Cy5 ^61^, 5’-GCAGCCACCCGAGGTGT-3’-Cy5 ^62^, 5’-GCTGCCACCCGTAGGTGT-3’-Cy5 ^62^, 5’-CCGCAACACCTAGTACTCATC-3’-Cy3 (targeting bacteria) ^63^.

Probe concentrations were 5 pmol μL^-1^ for Cy3 and Cy5, 8.3 pmol μL^-1^. Slides were incubated at 46°C for 1.5 h in humidity chambers equilibrated with hybridization buffer. Samples were washed using washing buffer containing per 50 mL 700 μL 5M NaCl, 500 μL EDTA, 1 mL 1M Tris/HCl pH 8.0, by adding and removing washing buffer (at 48 °C) several times after which the samples were incubated for 10 min with washing buffer by letting the samples float on top of a styrofoam Eppendorf holder in a 48°C water bath. The slides were held on ice and washed with ice cold distilled water in the same way as for the washing buffer and subsequently dried using compressed air. Samples were covered with "High Precision" No. 1.5H borosilicate coverslips (Marienfeld-Superior, Lauda-Königshofen, Germany) and visualized using the Leica SP8X confocal point scanning microscope equipped with hybrid HyD detectors and pulsed white-light laser. Observations were based on five biological replicates, two from experiment 3 and three from experiment 1.

### Scanning electron microscopy

Pieces of the anoxically fixed gold mesh film were subjected to an additional fixation with 2% paraformaldehyde and 2.5% glutaraldehyde in 0.1M PHEM (60mM Pipes, 25mM Hepes, 10mM EGTA, 2mM MgCl) buffer pH 6.9 for 1.5h on ice followed by three washes with PHEM Buffer. Samples were contrasted using 1% OsO_4_ with 1.5% K_3_[Fe(CN_6_)] for 1 hour on ice followed by four washes with ultrapure water. Dehydration was carried out using a graded acetone series followed by two washes with 100% ethanol and stepwise infiltration with hexamethyldisilazane (HMDS)^64^. Excess HMDS was removed by draining on filter paper. The gold mesh films were air dried overnight before mounting on specimen stubs using carbon tape. Samples were sputter-coated with palladium-gold prior to imaging using a JEOL 6330 Field emission SEM operating at 13kV using the in-lens detector.

### Transmission electron microscopy

Samples were processed as described above with the exception that after the dehydration the samples were stepwise infiltrated with EPON resin (2h 2:1, overnight 1:1, 3h 1:2, 3h 1:3 acetone:EPON, overnight 100% EPON) followed by embedding in freshly prepared EPON resin. After curing for 48h at 60°C the resin blocks were trimmed by hand using a razor blade. Ultrathin sections (50nm) sectioned on a diamond knife (Diatome ultra 45) using a Reichert-Jung ultracut E ultramicrotome and mounted on copper grids (100# hexagonal) with a carbon coated formvar film and post-stained using 2% uranyl acetate and Reynolds lead citrate before imaging in a JEOL 1400- Flash TEM operating at 120kV.

## Acknowledgements

This study was supported by the SIAM Gravitation grant funded by NWO [Grant number 024.002.002]. MJ was furthermore supported by the ERC Synergy Grant MARIX [Grant number 854088].

## Author contributions

HTO, RM, TS, MSMJ, AH, TB, CUW designed the experiments, HTO performed the experiments, MW performed the experiment using antibiotics, RM prepared the SEM and TEM samples, analysed them using the electron microscope and interpreted the data together with HTO and CUW, JP analysed the FISH samples using the confocal microscope together with HTO and interpreted the data together with HTO and CUW.

## Supplementary material

**Figure S1:**
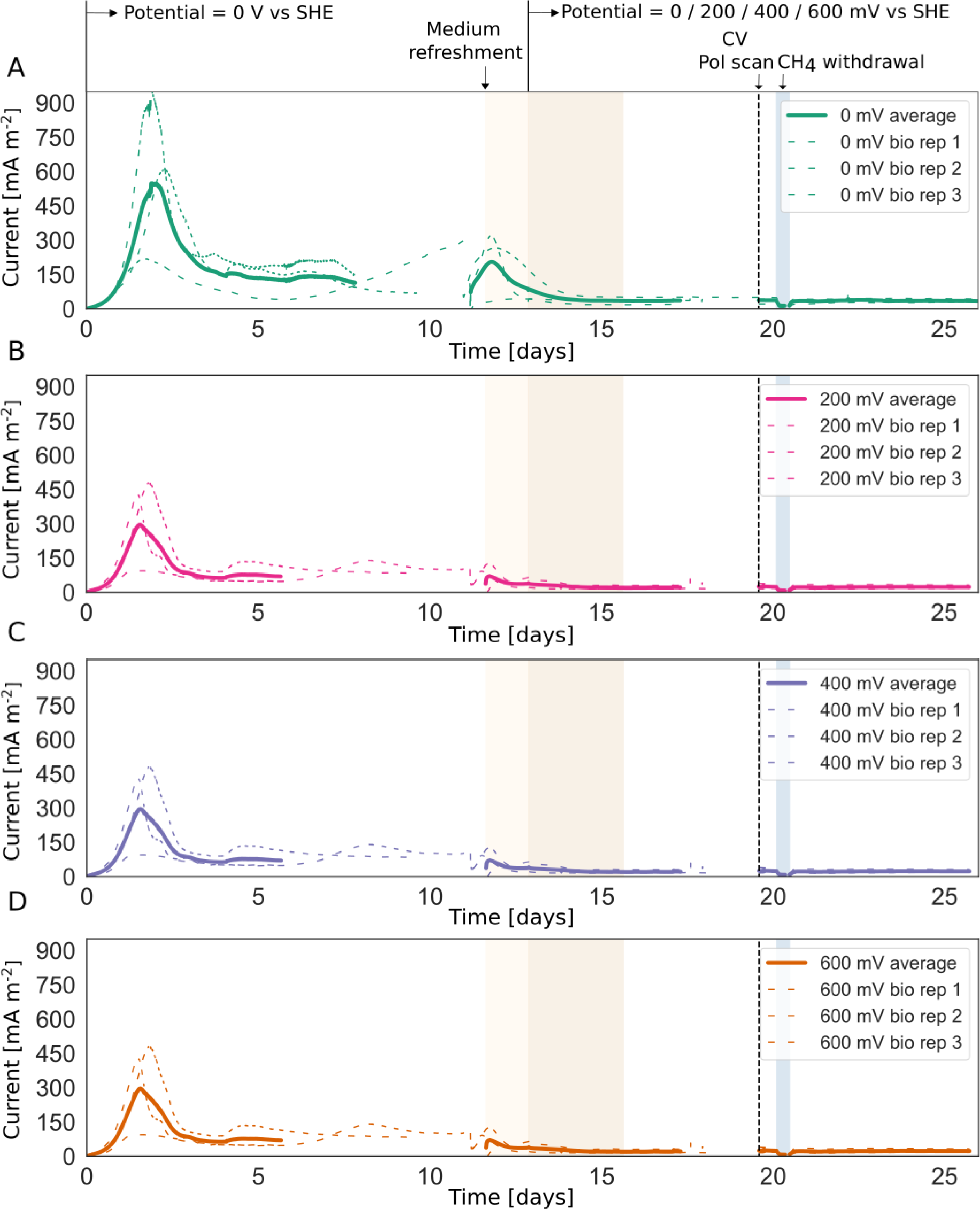
Bioelectrochemical data collected during experiment 1. Two distinct phases are visible: (i) a start-up phase during which all systems were operated at 0 mV versus the standard hydrogen electrode (SHE), represented by the first blank part and the light yellow part of the figure, with the beginning of the light yellow part indicating the time when the medium was refreshed, and (ii) a subsequent phase during which the potential was maintained at 0 mV or switched to 200 mV, 400 mV, or 600 mV vs SHE. In this part we conducted cyclic voltammetry scans and polarisation scans, as indicated by dashed line, and tested for methane-dependent current, depicted by the blue inset. At the end of the experiment, samples were collected for metagenome and metatranscriptome sequencing and the biofilm was visualized using microscopy.

**Figure S2:**
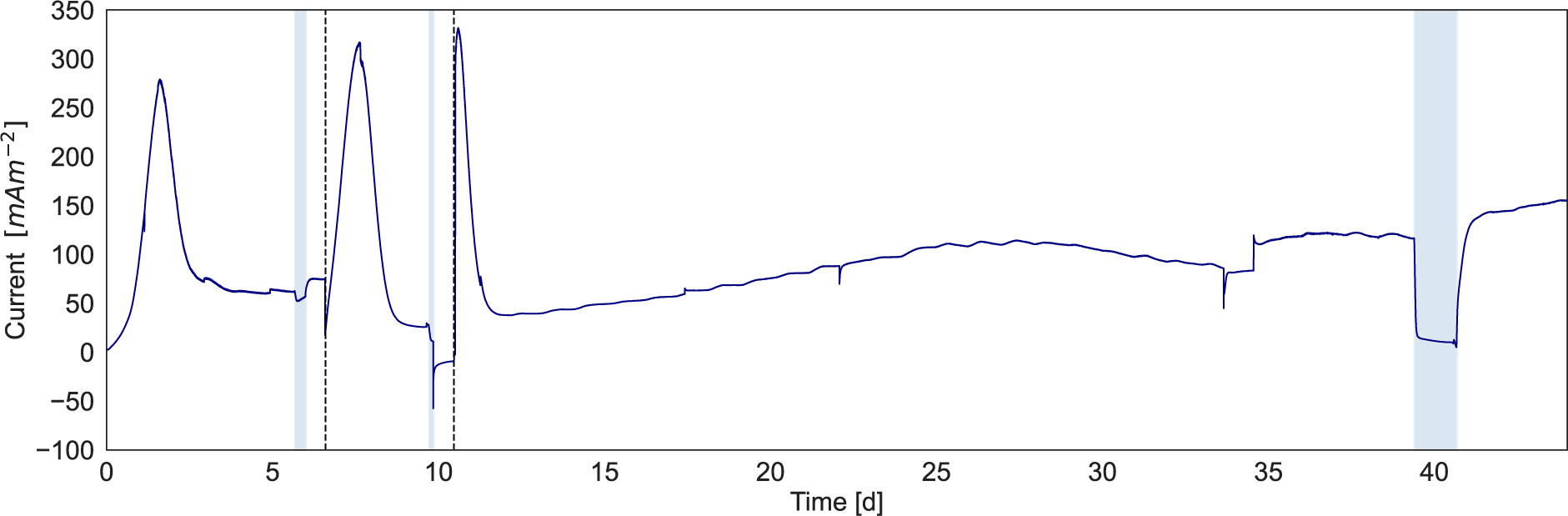
Bioelectrochemical data collected during experiment 2. The microbial community was incubated for 6.5 weeks at 0 mV vs SHE with two medium refreshments as indicated by the dashed lines. Methane-dependent current was measured as indicated by the blue insets. At the end of the experiment, samples were collected for metagenome sequencing.

**Figure S3:**
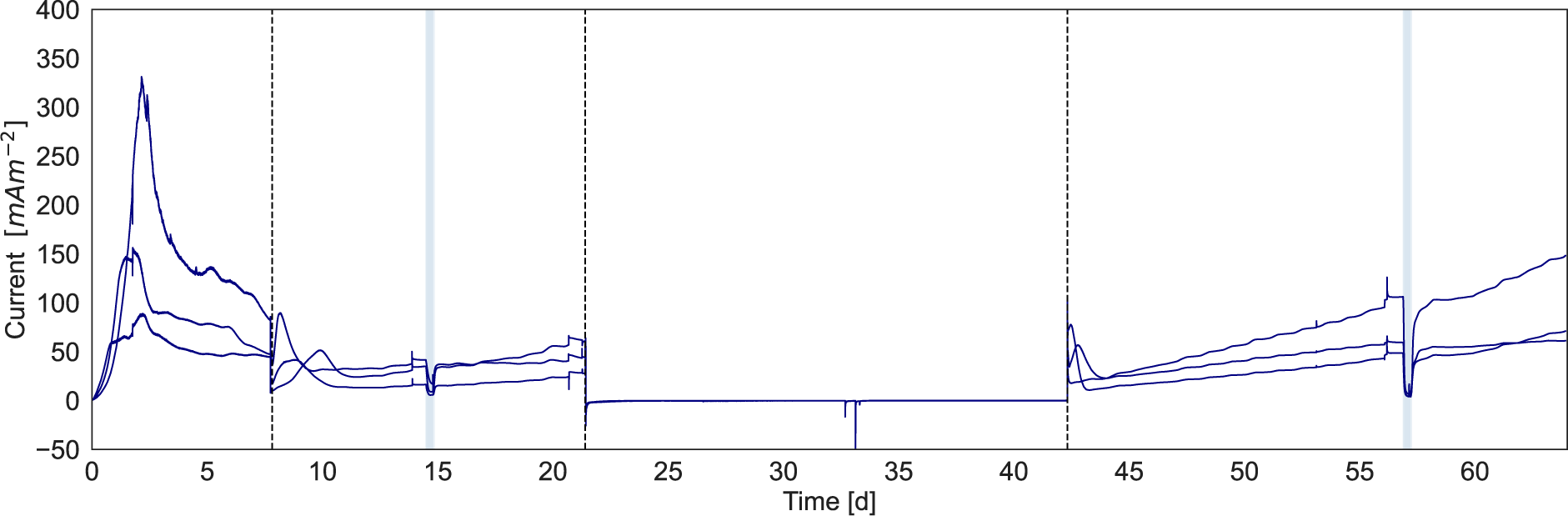
Bioelectrochemical data collected during experiment 3. The microbial community was incubated for 9 weeks at 0 mV vs SHE with three medium refreshments and a three-week famine phase at -400 mV vs SHE occurring between the second and third medium refreshment. The dashed lines represent the medium refreshments, and the famine phase is indicated by the dashed lines. Methane-dependent current was measured before and after the famine phase as indicated by the blue insets of the figure. At the end of the experiment, the biofilm was visualized using microscopy.

**Figure S4:**
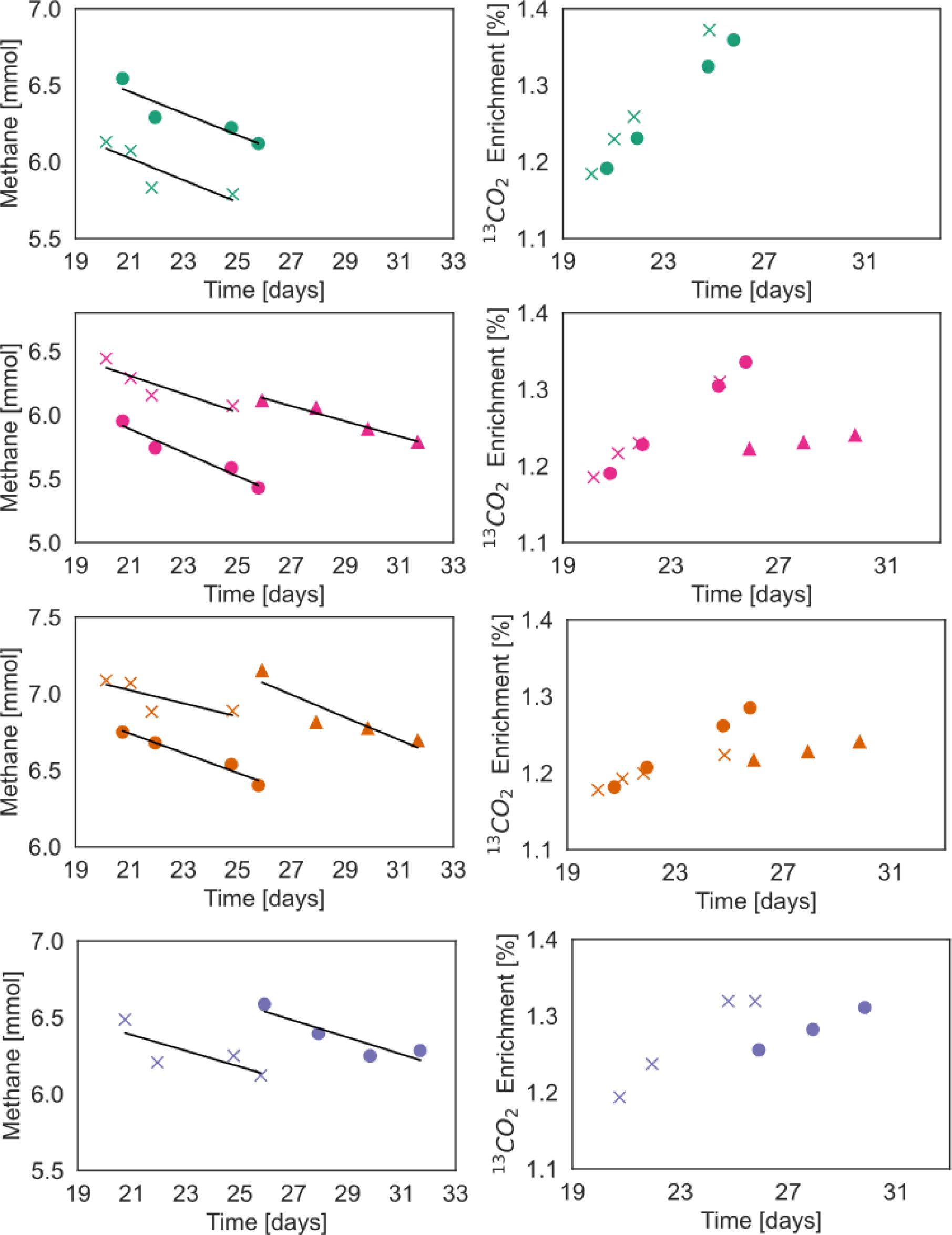
Total methane consumption (left) and relative enrichment of ^13^CO_2_ (expressed as a percentage) compared to total CO_2_ (right) for experiment 1. Colours correspond to conditions reported in Figure S1: green represents 0 mV, pink represents 200 mV, orange represents 400 mV and purple represents 600 mV. These results were collected at the end of the experiment after introducing ^13^CH_4_ to a sealed system. From this experiment the Coulombic efficiency was calculated to be 5.5% ± 0.013, which was calculated using formula 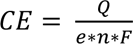 with CE coulombic efficiency, Q the cumulative charge expressed in Coulombs (C), the moles of electrons per mole methane, n the moles of methane being consumed and F the Faraday constant being 96485 C mol^-1^. However, it is challenging to draw conclusions from this measurement due to the large number of samples that was taken over a short period of time increasing the chances for methane to escape and making the calculation inaccurate.

**Figure S5:**
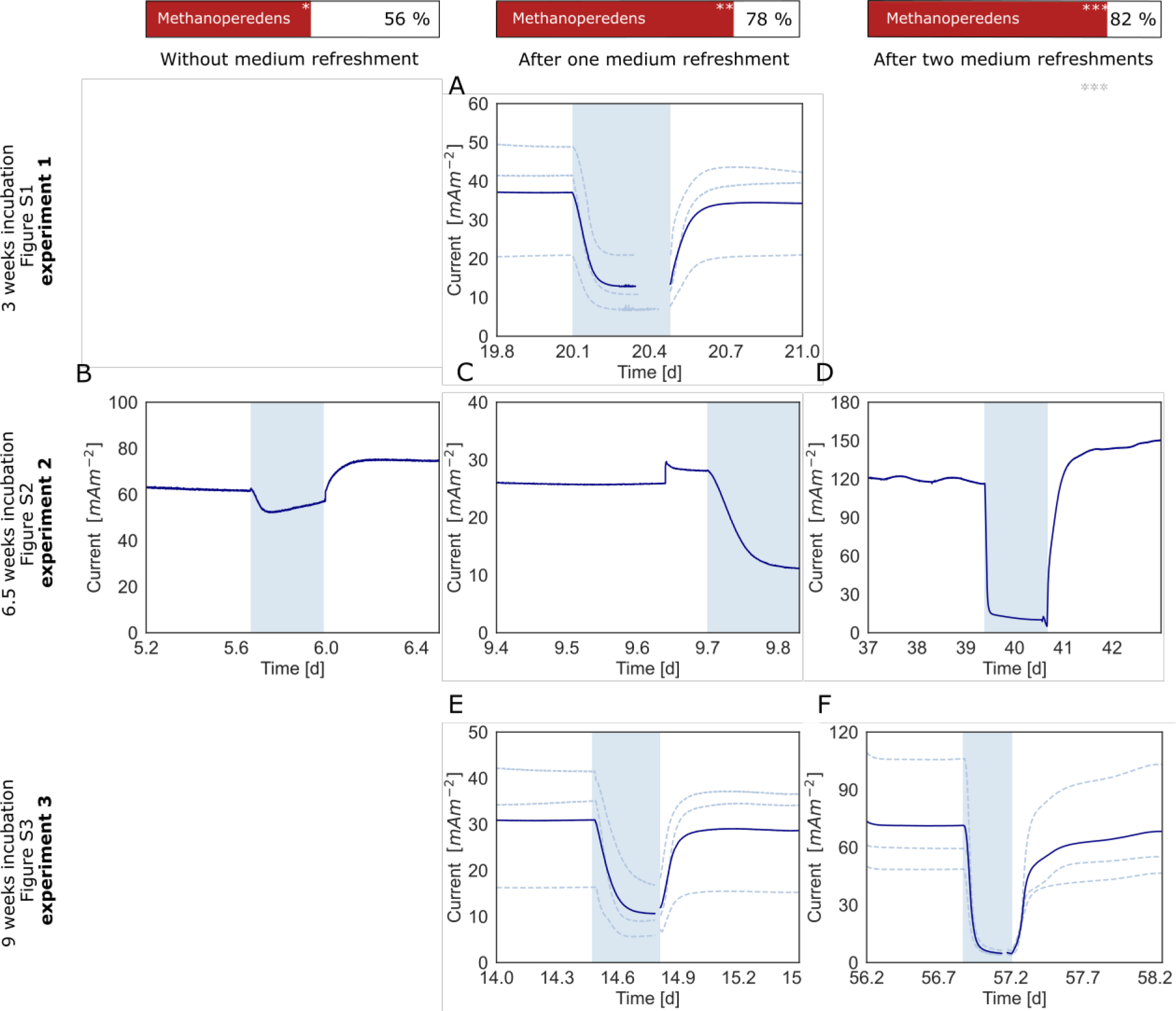
Influence of the number of medium refreshments accompanied by a longer incubation time on the methane-dependent current and relative abundance of ‘*Ca*. Methanoperedens’. This Figure shows the primary data that is related to Figure 3.

**Figure S6:**
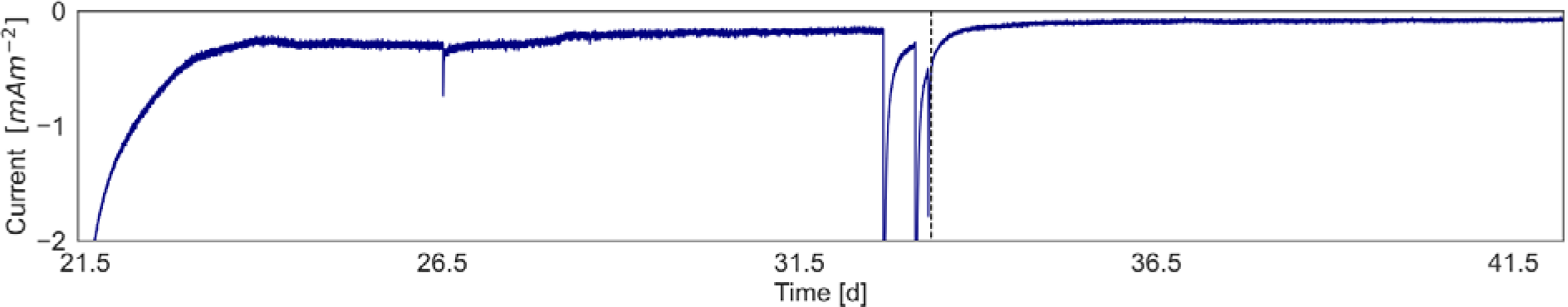
Bioelectrochemical data during the famine phase in which the potential was -400 mV vs Standard Hydrogen Electrode (SHE). At 33.4 days, the system was closed to measure the production of methane in batch as depicted by the dashed line.

**Figure S7:**
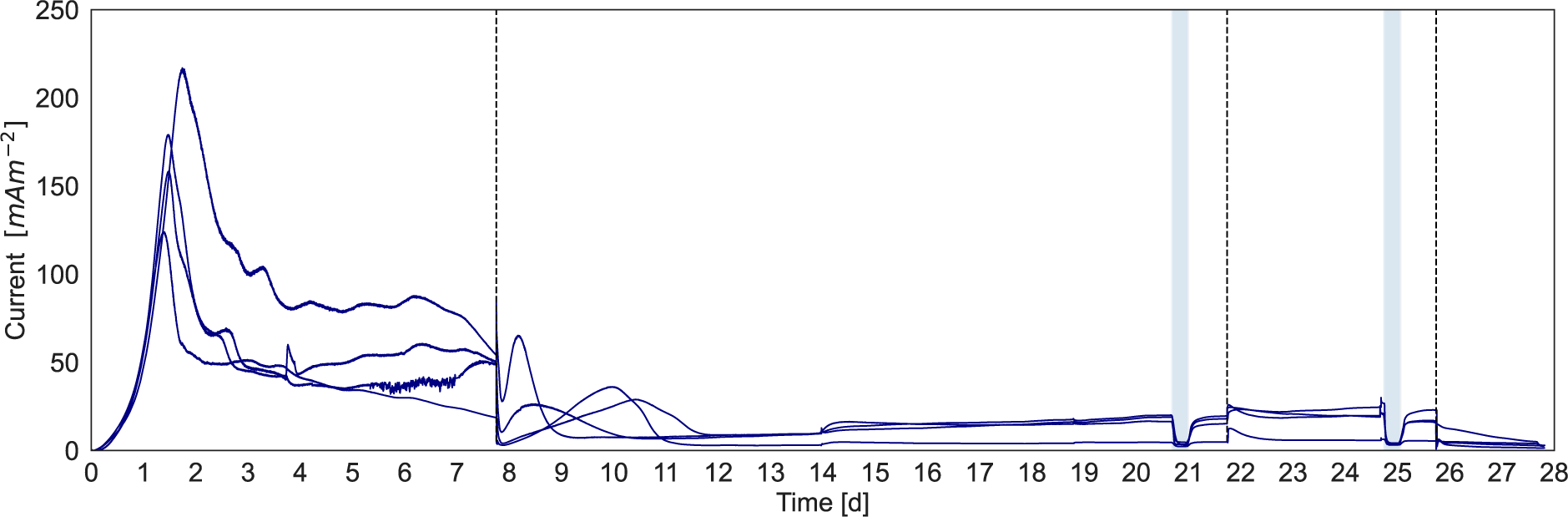
Bioelectrochemical data collected during the experiment in which we tested the influence of a bacterial antibiotic mixture (vancomycin, streptomycin, ampicillin and kanamycin at concentrations of 50 μg mL^-1^ each). Four biological replicates were run. Before and after the addition of antibiotics the methane-dependent current was tested as depicted by the blue insets. Near the end of the experiment, depicted by the third dashed line, we tested the influence of 2- bromoethanesulfonate (BES) (20 mM) in two biological replicates (Figure S8) and the influence of the archaeal antibiotic puromycin (50 μg mL^-1^) affecting the production of ribosomes in two other biological replicates (Figure S9). Medium refreshment is depicted by the first dashed line.

**Figure S8:**
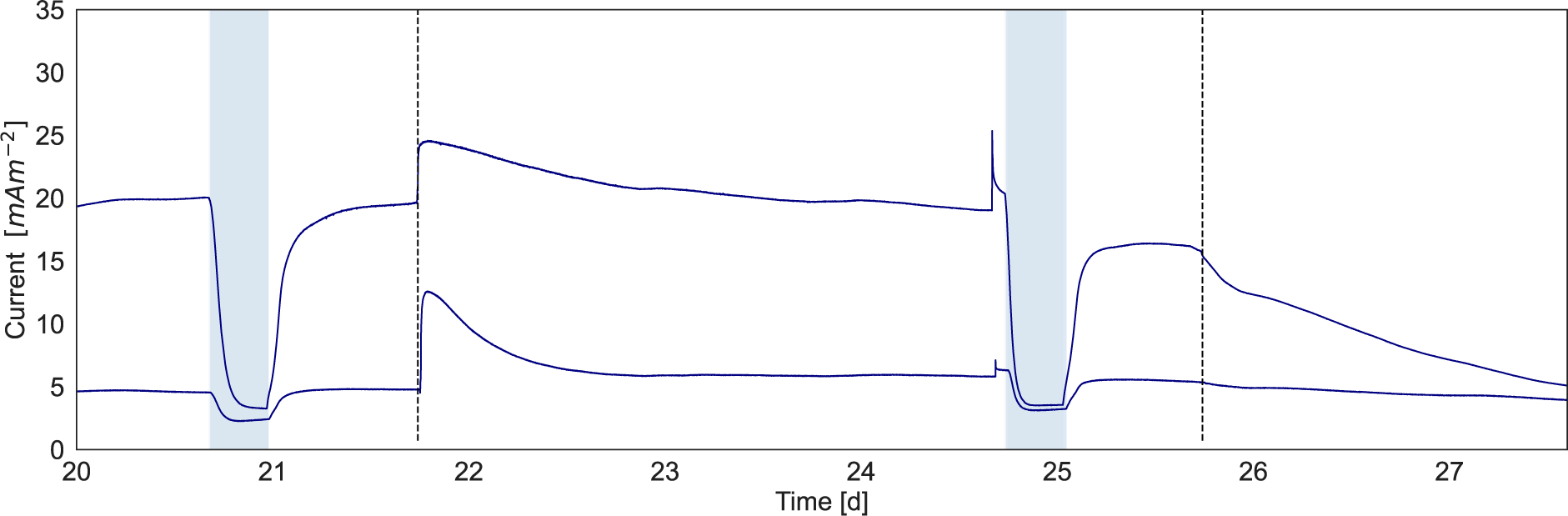
Bioelectrochemical data collected during the experiment in which the influence of antibiotics was tested; enlarged from Figure S7. The first dashed line indicates the addition of the bacterial antibiotics mixture and the second dashed line indicating the addition of archaeal antibiotic puromycin (50 μg mL^-1^). The blue inset indicates where argon was used in the gas phase instead of methane.

**Figure S9:**
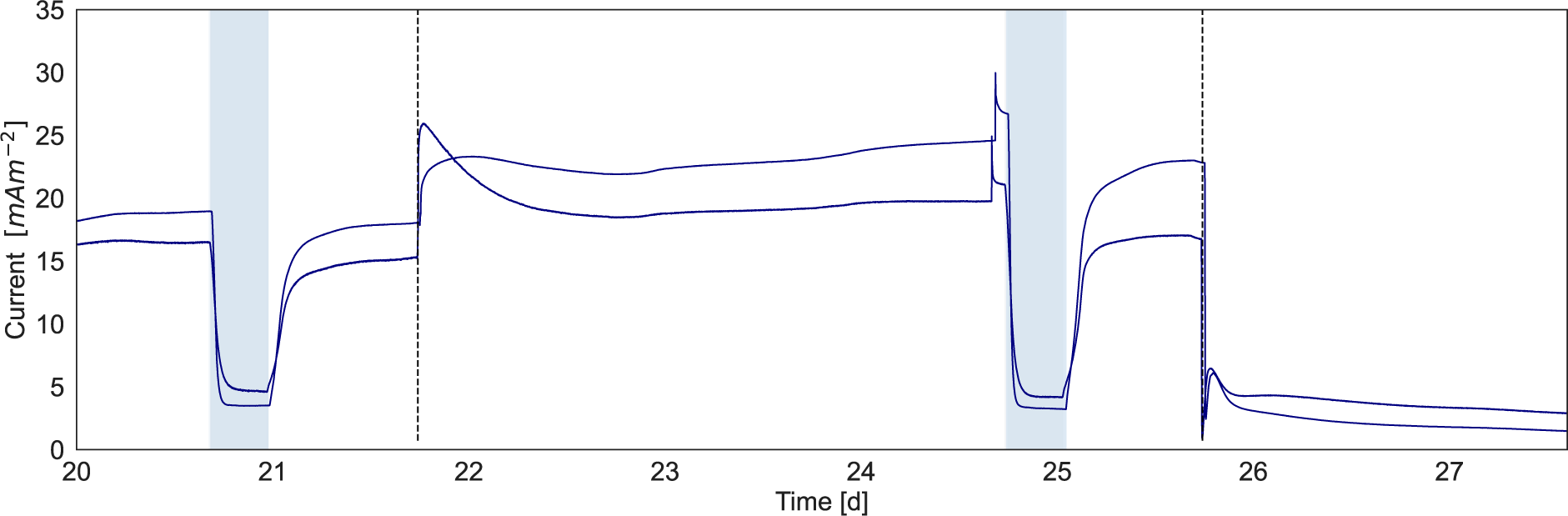
Bioelectrochemical data collected during the experiment in which the influence of antibiotics was tested, enlarged from Figure S7. The first dashed line indicates the addition of the bacterial antibiotics mixture and the second dashed line indicating the addition of MCR inhibitor 2- bromoethanesulfonate (20 mM). The blue part indicates the part where argon was used in the gas phase instead of methane.

**Figure S10:**
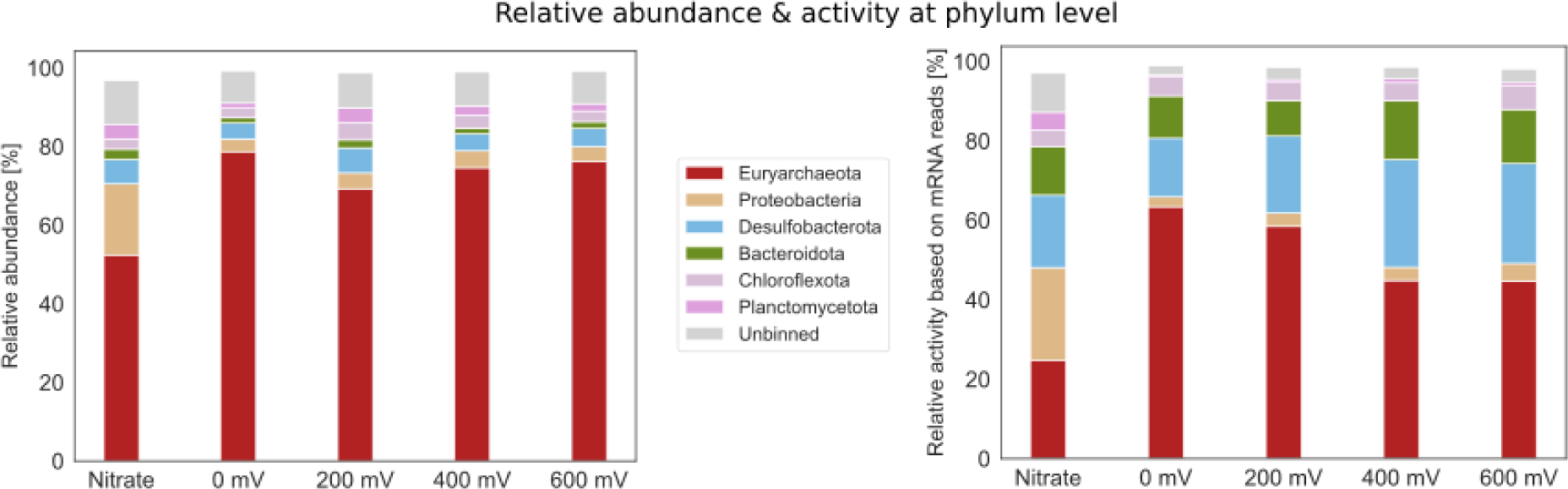
Relative abundance and relative activity at phylum level determined by mapping nucleic acid and mRNA reads to the metagenome assembled genomes (MAGs). The Methanoperedenaceae and Geobacteraceae family both contain only one MAG that was identified as ‘*Ca*. Methanoperedens’ and *Geobacter* sp., respectively, based on the classification by the Genome Taxonomy Database (GTDB) in this study. The completeness of the ‘*Ca*. Methanoperedens’, *Geobacter* sp. and Ignavibacteriaceae bin is 99.3%, 99.4% and 97.2% and the contamination of these bins is 5.2%, 0% and 0.6. For these figures we applied a threshold of 1% relative abundance.

**Table S1:** Relative abundance and relative activity of all community members classified using GTDB-Tk

**Table S2:** Overview of all metatranscriptomics data

